# A new microphysiological system shows hypoxia primes human ISCs for interleukin-dependent rescue of stem cell activity

**DOI:** 10.1101/2023.01.31.524747

**Authors:** Kristina R. Rivera, R. Jarrett Bliton, Joseph Burclaff, Michael J. Czerwinski, Jintong Liu, Jessica M. Trueblood, Caroline M. Hinesley, Keith A Breau, Shlok Joshi, Vladimir A. Pozdin, Ming Yao, Amanda L. Ziegler, Anthony T. Blikslager, Michael A. Daniele, Scott T. Magness

## Abstract

**Background & Aims:** Hypoxia in the intestinal epithelium can be caused by acute ischemic events or conditions like Inflammatory Bowel Disease (IBD) where immune cell infiltration produces ‘inflammatory hypoxia’, a chronic condition that starves the mucosa of oxygen. Epithelial regeneration after ischemia and IBD suggests intestinal stem cells (ISCs) are highly tolerant to acute and chronic hypoxia; however, the impact of acute and chronic hypoxia on human ISC (hISC) properties have not been reported. Here we present a new microphysiological system (MPS) to investigate how hypoxia affects hISCs isolated from healthy human tissues. We then test the hypothesis that some inflammation-associated interleukins protect hISCs during prolonged hypoxia.

**Methods:** hISCs were exposed to <1.0% oxygen in the MPS for 6-, 24-, 48- & 72hrs. Viability, HIF1α response, transcriptomics, cell cycle dynamics, and hISC response to cytokines were evaluated.

**Results:** The novel MPS enables precise, real-time control and monitoring of oxygen levels at the cell surface. Under hypoxia, hISCs remain viable until 72hrs and exhibit peak HIF1α at 24hrs. hISCs lose stem cell activity at 24hrs that recovers at 48hrs of hypoxia. Hypoxia increases the proportion of hISCs in G1 and regulates hISC capacity to respond to multiple inflammatory signals. Hypoxia induces hISCs to upregulate many interleukin receptors and hISCs demonstrate hypoxia-dependent cell cycle regulation and increased organoid forming efficiency when treated with specific interleukins

**Conclusions:** Hypoxia primes hISCs to respond differently to interleukins than hISCs in normoxia through a transcriptional response. hISCs slow cell cycle progression and increase hISC activity when treated with hypoxia and specific interleukins. These findings have important implications for epithelial regeneration in the gut during inflammatory events.

## INTRODUCTION

Cellular respiration in complex multi-layered tissues relies on a constant supply of oxygen to maintain healthy physiologic states. To this end, the intestinal epithelium maintains physiologic oxygen tension through a highly vascularized tissue that is perfused by a vast network of vessels that terminate in capillary beds where oxygen is released to adjacent cells in a local microenvironment.^1^ In the crypt-villus epithelial architecture, cells can experience marked differences in physiologic oxygen concentrations where crypt-based cells experience higher oxygen levels compared to cells at the villus tips.^2^ Along the crypt-villus axis, a steep oxygen gradient is present wherein microenvironments that are separated by just ∼100 cell distances experience a ∼10-fold lower oxygen concentration.^3^ While lower oxygen concentrations within the gradient are tolerated as normal, sudden or dramatic changes in the magnitude and duration of oxygen loss can lead to pathological hypoxia resulting in cell dysfunction and/or death.^4^

Most studies that evaluate the impact of tissue oxygen deprivation do so in the context of ischemia-reperfusion injury. This is characterized by an initial ischemic event such as bowel strangulation, aneurysm, or organ harvest for transplantation which induces hypoxia. Subsequent reperfusion of blood flow, and thus oxygen, induces a separate type of oxidative injury caused by free radicals and reactive oxygen species.^5,6^ Hypoxic events can be categorized as acute, which are typically transient events that last on the order of minutes to hours, and chronic hypoxic events that last much longer, for at least 24 hours or more.^7^ Inflammatory hypoxia, a type of chronic hypoxia, is caused by the presence of large numbers of immune cells in the submucosal compartment depleting local oxygen supplies.^4,8,9^ Following epithelial injury, a massive influx of immune cells is recruited to prevent pathogens and other luminal contents from causing further damage. This massive influx of immune cells consumes the local oxygen supply to the epithelium^4,10-13^ and can cause vasoconstriction, which further starves the epithelial cells of oxygen.^3,14,15^ Hypoxia due to inflammation can exist in isolated microenvironments that might initially go unnoticed and eventually resolve themselves. However, longer periods of inflammation, such as in Inflammatory Bowel Disease (IBD) may compromise the epithelium and normal repair mechanisms ultimately resulting in, or contributing to, mucosal erosions and ulcerations.^8,12^ Consistent with this concept, inflammatory responses have been shown to produce localized and chronic hypoxic episodes in the intestinal stem cell (ISC) zone, potentially leading to impaired ISC survival, renewal, and differentiation, ultimately impeding mucosal repair and producing ISC death.^6,15,16^

While numerous studies focus on the impact of hypoxia on repair mechanisms involving restitution in differentiated entrocytes^6,17-21^ few studies investigate the impact of hypoxia on ISC properties and ISC-dependent repair mechanisms^22^, especially in human tissues. Traditionally, human cancer cell lines are used to study acute and chronic hypoxia at the cellular level but transformed cells exhibit abnormal proliferation and cell death dynamics and they do not properly differentiate, thus they are poor models of human ISCs (hISCs).^23^ Studies in porcine small intestines show that the so-called ‘reserve’ ISC (rISC) exhibits high tolerance to hypoxic episodes of up to 4 hours of hypoxia/ischemia^15,22^, but these studies do not evaluate the impact of chronic hypoxia on hISCs. A highly physiologically relevant human model to study acute ischemia was developed^24^ and while the study primarily characterizes the differentiated epithelium under hypoxia, a lack of apoptosis in the crypt observed after 45 min of ischemia indicates hISCs are highly resistant to short hypoxic episodes.^20^ Despite the important observations and advances outlined in these in vivo studies, all have been performed either in the context of ischemia-reperfusion injury, precluding the evaluation of hypoxic injury uncoupled from reperfusion injury, or performed under short-duration ischemic conditions (<4 hours), and thus interpretations are limited to acute rather than chronic hypoxia injury.

Investigating the impact of hypoxia on hISCs has been historically limited by the inability to indefinitely culture hISCs, the lack of high-resolution in vitro platforms where oxygen levels can be tuned and accurately monitored in real time, and ethical considerations of human research that preclude in vivo experimentation. In this study, we engineered a microphysiological system (MPS) that delivers multiple durations and magnitudes of hypoxia to primary hISCs. The MPS allows for precise remote monitoring and control of oxygen concentrations in real-time at the cell surface interface. We use the MPS to create controlled durations of acute and chronic hypoxic episodes and evaluate dynamic transcriptomic changes within hISCs under these conditions. We focused our evaluation on chronic hypoxic events related to hISC-immune cell cross talk through interleukin signaling that might occur during inflammatory hypoxia. Our findings have broad implications for understanding the role in which inflammatory hypoxia can impact hISC behavior in gross clinical presentation of ischemia-reperfusion injury and focal ischemic events that occur in Inflammatory Bowel Disease-related hypoxia.

## RESULTS

### A new microscopy-compatible microphysiological system enables precise regulation and real-time monitoring of oxygen levels at the cell-media interface

To address the limitations of sensitivity, accuracy and scale in existing environmental culture systems that create a hypoxic environment, we developed a novel MPS the size of a standard microscope slide where oxygen tension could be tailored to precise levels. The MPS design consists of 2 major structural components, 1) the 3D-printed light- and air-tight container required for phosphorescence detection and hypoxia induction in the closed system and 2) a clear-bottomed acrylic five-well tissue culture plate with each well containing a hydrogel scaffold that facilitates hISC expansion, which all sits inside of the sealed chamber^25^ (**Fig. 1A,B**, see methods).

**Figure 1.**
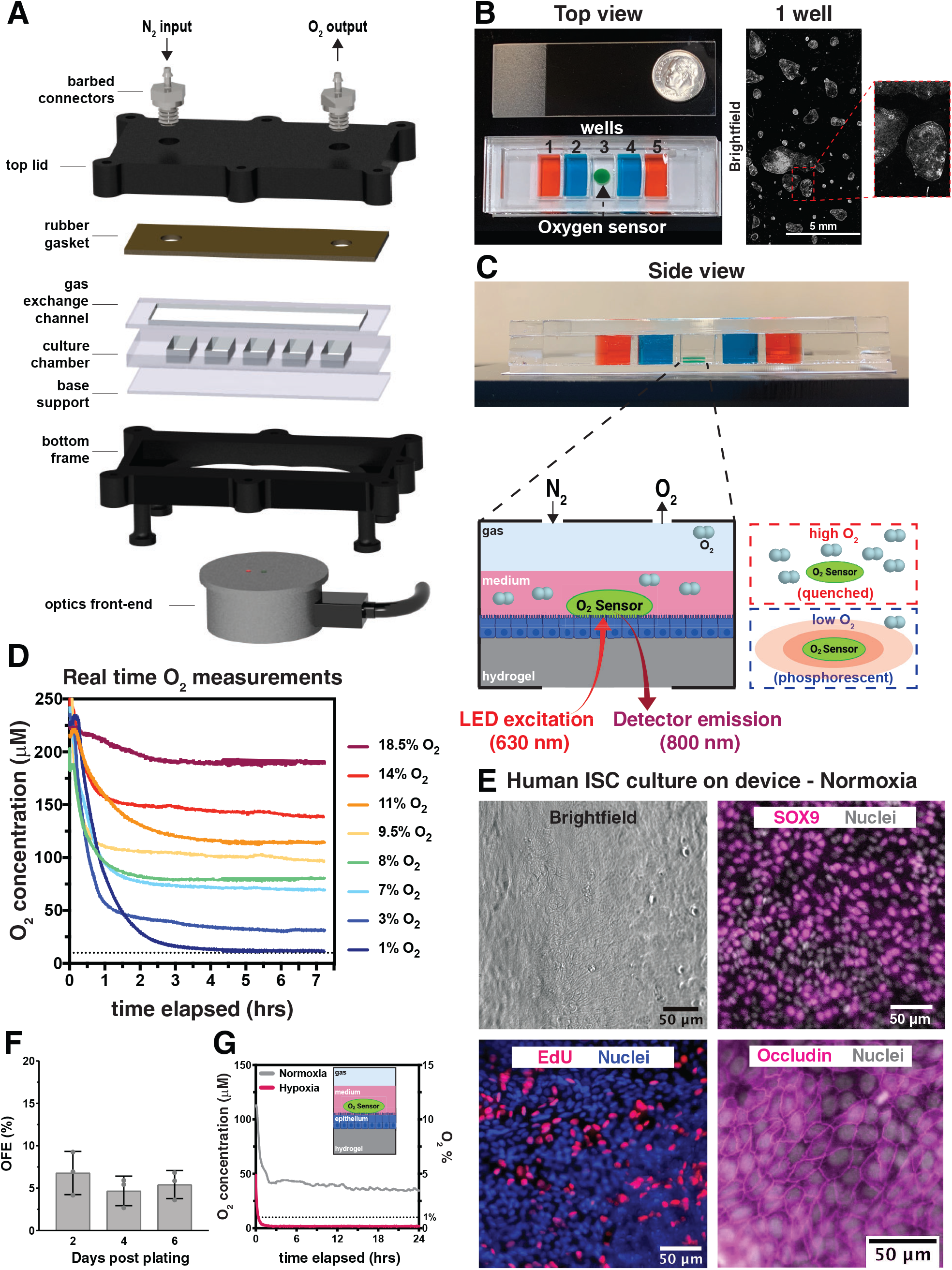
Development of a tunable hypoxic microphysiological system (MPS) with integrated oxygen sensors and cultured primary human intestinal epithelium. **A**) Exploded view of MPS compartments. **B**) (left) Top view photograph of a standard microscope glass slide (75 × 25 mm) and dime, for reference, placed above the MPS with iPOB (black arrow) inside well 3 and red and blue dyes in wells 1 and 5 and 2 and 4, respectively. (middle) 1 well brightfield image of monolayer of cells. (right) Inset from 1 well brightfield image shows magnified colonies in hISC monolayer. **C**) (Top) Side view photograph of MPS with middle well containing the green iPOB. Schematic of iPOB integrated into hydrogel to measure oxygen at the cell layer inside the cell culture well. (Bottom-left) Schematic of porphyrin-based luminescence. In high oxygen, luminescence from the porphyrin is quenched by energy transfer to oxygen, resulting in a decrease in phosphorescence lifetime. In the absence of oxygen, porphyrin molecules are excited by LED from the detector and phosphoresce with increased lifetime. **D**) Plots of 8 oxygen concentration versus time (7.5 hours) measurements from the iPOB inside the MPS, with 8 different mixed gas inputs, show generation of 8 established oxygen environments. **E**) (Top-left) Brightfield image of human intestinal epithelial stem and progenitor cells grown inside a MPS well to confluence. Fluorescent images show tight junction structures between epithelial cells marked by the protein Occludin (magenta, bottom-right), stem cells marked by SOX9 (magenta, top-right, and proliferative cells marked by EdU (red, bottom). Nuclei are marked by Bisbenzimide+ staining in grey or blue. **F**) Organoid forming efficiency (%) measured over 6 days from single cells isolated from MPS after 5 days in culture. **G**) Oxygen concentration tracked using the iPOB inside a MPS containing human intestinal epithelium for 24 hours with normoxic and hypoxic culture environments.

The MPS was designed as a closed system in which ambient air flowed over each well via continuous delivery of mixed gases. A biocompatible integrated Phosphorescent Oxygen Biosensor (iPOB) was placed on the hydrogel scaffold surface to provide real-time oxygen measurements specifically at the cell surface interface (**Fig. 1C**)^26^. Oxygen levels were quantified by the iPOB using Near Infrared (NIR) phosphorescence lifetime fluorimetry which measures the oxygen-dependent decrease in phosphorescence lifetime to determine local oxygen concentration (**Fig. 1C,D**)^27^. The iPOBs are reusable, highly sensitive, and capable of detecting a broad range of oxygen levels created by an off-chip gas mixer^28^ (**Fig. 1D**, see methods).

The MPS design facilitates easy access to refresh media, add compounds, and perform a variety of downstream assays including microscopy, transcriptomic analysis, and immunostaining. The MPS is fabricated from relatively inexpensive commercially available materials and can be sterilized and reused many times. Notably, multiple MPSs can be connected to the same gas source to accommodate complex experimental designs that require many perturbations and technical replicates (see methods). Together, these demonstrate an ideal model system for evaluating the effect of hypoxia on hISCs.

### The MPS supports culture of hISC/progenitor epithelial monolayers

We recently developed methods that support indefinite 2-Dimensional (2D) expansion of proliferative human intestinal stem cells from primary small intestine and colon^25^. Under normoxic conditions (i.e., 37°C, 5% CO_2_, and remainder atmospheric air), hISC monolayers can be cultured on a defined hydrogel and media formulation that supports hISC expansion while repressing differentiation^25,29^. This method was used to expand primary jejunal hISCs isolated from small intestinal crypts of an organ transplant donor and then transferred to the MPS. Initial characterization of the proliferative hISC monolayer was performed in the MPS under normoxic conditions by immunostaining. The monolayers demonstrated DNA synthesis as measured by S-phase marker EdU, expression of hISC/progenitor cell marker SOX9^*30,31*^, and epithelial cell tight junction protein Occludin (OCLN) **(Fig. 1E)**. These results demonstrate the MPS is biocompatible with primary hISCs that are consistent with proliferating hISC/progenitor cell populations.

Next, the baseline functional stemness of proliferative hISCs cultured on the MPS under normoxia was evaluated. hISCs were cultured on the MPS as proliferative monolayers under atmospheric O_2_ levels, dissociated to single cells and plated in Matrigel™ on a CellRaft™ Array to assess organoid forming efficiency (OFE), a measure for in vivo hISC activity^32-34^. Single cells in the microwells were identified, quantified, and putative hISCs were individually tracked over a 6-day culture period to determine whether an organoid formed. The data demonstrate ∼4% of single cells cultured in the MPS under hISC expansion conditions generated an organoid at each time point measured (2, 4, and 6 days after plating) **(Fig. 1F)**, which is consistent with OFE of FACS-isolated hISCs from a Lgr5-EGFP expressing reporter gene mouse and also consistent with human organoid-derived hISCs cultured for the same period of time in synthetic matrices^34,35^. These findings demonstrate that hISCs cultured as 2D monolayers on the MPS maintain hISC properties over time, and at similar ratios to those observed in vivo and in 3D organoid systems.

To define the speed at which severe hypoxia could be achieved and maintained in the MPS, an iPOB was added to the epithelial cell surface, and real-time oxygen levels at the cell-media interface were measured for 24 hours **(Fig. 1G**, *inset*). Oxygen concentrations of <1% (below 10 μM) are considered severely hypoxic but not anoxic^36^. The data show the MPS produced a rapid hypoxic environment of 0.3% O_2_ (3 μM) within ∼30 minutes, which was constantly maintained during the 24-hour experiment **(Fig. 1G)**. Interestingly, when proliferative hISC monolayers were cultured in normoxic conditions as a control, cellular respiration reduced oxygen levels at the cell-media interface from 18% (180 μM) to 3.5% (35 μM) in the MPS. These data demonstrate that physiologic O_2_ metabolism of proliferative hISCs produce a steady-state flux of O_2_ in just 30-minutes that is 5-fold less than atmospheric oxygen levels.

### Human ISC monolayer cultures demonstrate peak HIF1α response at 24 hours of severe hypoxia with no loss of stem cell activity after 48 hours of severe hypoxia

The HIF1α transcription factor^13^ is a master regulator of the hypoxic response conserved across all tissues and species.^14,37^ Under normoxia, HIF1α is constitutively expressed but is immediately targeted for degradation^14^. By contrast, under hypoxia, HIF1α degradation is inhibited through well-established mechanisms, resulting in rapid accumulation and regulation of a broad range of downstream target genes that regulate cell survival, proliferation, metabolism, and cell migration^13,38^. Immunostaining for HIF1α in hISCs under durations of hypoxia shows a significant accumulation of HIF1α only at 24 hours of severe hypoxia **(Fig. 2A,B)**. No significant change in HIF1α was observed at 6 hours of hypoxia compared to normoxia, which may indicate tolerance of proliferative hISCs to shorter hypoxic episodes. Lack of HIF1α accumulation at 48 hours of hypoxia is likely due to known dampening of the HIF-response that occurs after cells re-equilibrate to tolerate new and sustained low O_2_ levels.^4^ Interestingly, hISCs cultured under normoxia showed a significant increase in HIF1α levels 24 hours after media change when compared to the 6 hour normoxic samples. This minor hypoxic response is consistent with previous observations that cells in submersion cultures experience a localized hypoxic environment due to the slower rate of diffusion of O_2_ through medium relative to the rate of consumption in cellular metabolism.^39^ Together, these findings established a timeframe for further investigation into the mechanisms by which hISCs resist death and preserve functional stemness and viability in response to HIF1α accumulation in hISCs during short-, medium-, and long-term hypoxia.

**Figure 2.**
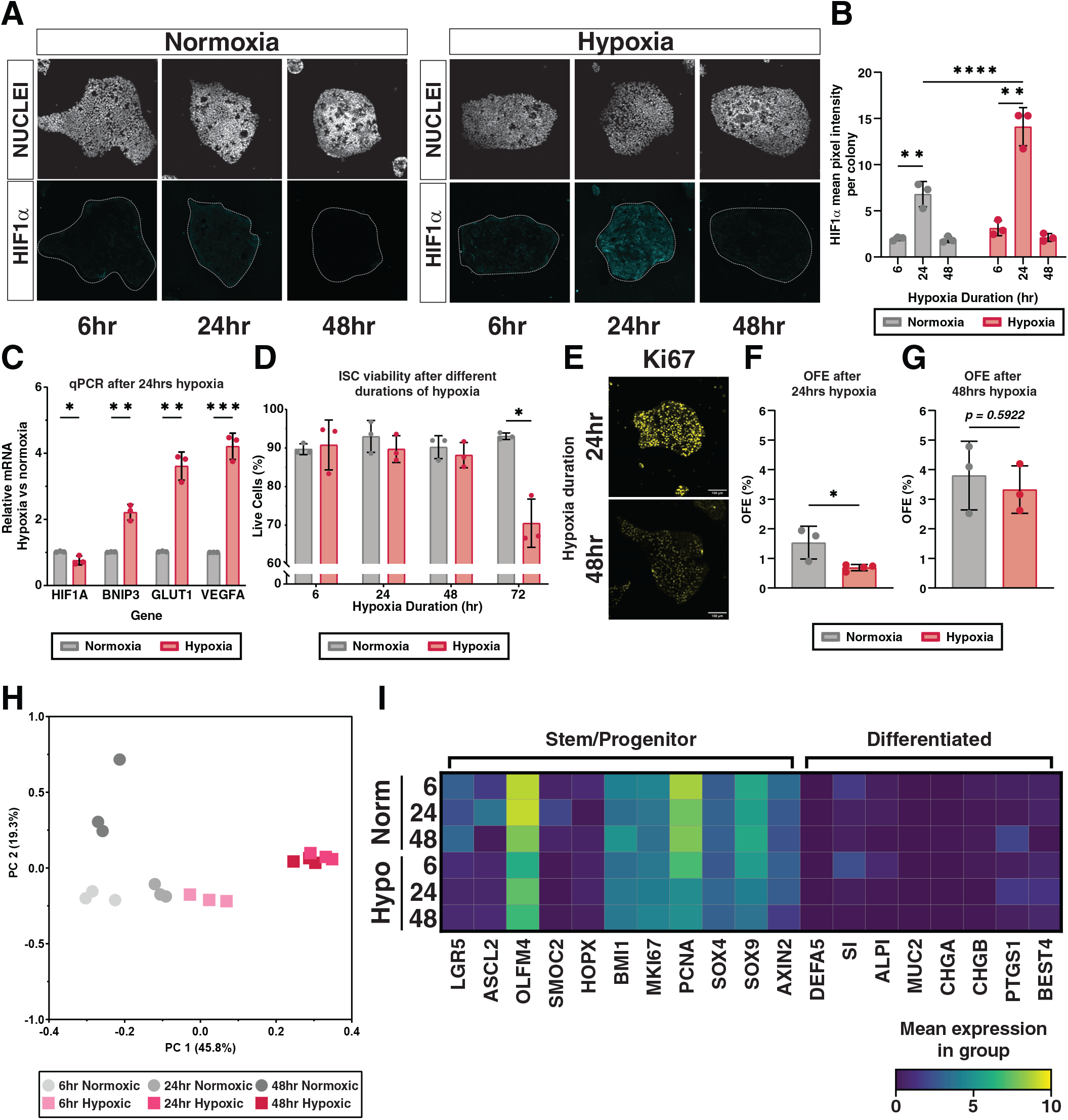
Primary human intestinal epithelium cultured under hypoxia inside microphysiological system shows dynamic response to different durations of hypoxia. **A**) Representative images showing colonies of human intestinal stem cells cultured on collagen scaffolds as 2D monolayers showing dynamic changes in HIF1α IF staining over different durations of normoxia or acute hypoxia. **B**) Mean pixel intensity of HIF1α staining in **A** over different durations of normoxia or hypoxia. Technical replicates represent mean pixel intensities of all nuclei in 3 individual colonies. **C**) RT-qPCR results of human intestinal epithelium for downstream HIF1α targets (BNIP3, GLUT1, VEGFA) after 24 hours of normoxic or hypoxic environment exposure. **D**) Flow cytometry results from single cells isolated after 6, 24, 48, and 72 hours from normoxic and hypoxic MPS and stained for Annexin V(-) to quantify cell viability cells (%)**E**) Immunostaining of hISC monolayer colonies for proliferation marker KI67 following 24 or 48 hours of hypoxia **F**) Functional stemness as measured by Organoid forming efficiency (OFE %) for 24-hour normoxic and hypoxic samples after 6-days of 3D Matrigel™-embedded culture. **G**) Functional stemness for 48-hour normoxic and hypoxic samples after 6-day of 3D Matrigel™-embedded culture. **H**) Principal component analysis of RNA-seq results showing clusters of replicate samples from each time point (6, 24, 48 hours) and condition (normoxia or hypoxia). Bulk RNA-sequencing samples were sequenced from a range of 39 to 93 million reads **I**) Heatmap showing mean CPM-normalized and log-transformed expression of stem/progenitor markers or differentiated lineage markers from hypoxia bulk RNA sequencing in **2H**.

To confirm whether proliferative hISCs in the MPS were experiencing a HIF-response at 0.3% O_2_, qPCR was performed to detect expression of canonical HIF1α target genes that are typically upregulated during hypoxia. After 24 hours of hypoxia, *HIF1α* demonstrated little transcriptional change as expected since its activity is mediated by post-transcriptional stabilization (**Fig 2C**)^40,41^. By contrast, the set of classic HIF1α target genes (*VEGF, SCL2A1/GLUT1, BCL2, BNIP3*) were all significantly upregulated compared to normoxic controls **(Fig. 2C)**^42-44^. These findings demonstrate that an O_2_ tension of 0.3% (3 μM) for 24 hours is sufficient to activate a canonical hypoxia response inside proliferative hISCs.

Next, we sought to explore the extent to which hISCs can survive severe hypoxia. hISCs were exposed to severe hypoxia (<1% O_2_ as measured by the iPOB) in the MPS for 6, 24, 48, or 72 hours and viability was evaluated by staining for early apoptosis marker, Annexin V (**Fig. 2D**).^45^ Flow cytometric quantification of Annexin V negative cells demonstrates ∼90% viability through 48 hours of severe hypoxia. However, at 72 hours of severe hypoxia, there was a significant decrease in viability as measured by Annexin V-negative staining, from ∼90% viable in normoxic controls to ∼70% viable after 72 hours of hypoxia.

To determine how longer episodes of hypoxia would impact hISC activity, monolayers were exposed to 24 hours or 48 hours of hypoxia or normoxia as a control in the MPS. At 24 and 48 hours of hypoxia, hISCs demonstrate persistent expression of the proliferation marker, KI67, albeit nuclear KI67 levels are reduced at 48 hours of hypoxia **(Fig 2E)**. To quantify hISC activity after these durations of hypoxia, monolayers were subjected to 24 or 48 hours hypoxia, dissociated to single cells, and then applied to CellRaft™ Arrays (CRAs) for high-throughput accurate quantification of clonal OFE.^34^ CRAs enable up to ∼10,000 hISCs to be plated in microwells on a small device and individual hISCs can be evaluated for OFE over time^34,46^. After 24 hours of hypoxia OFE decreased ∼2.5-fold **(Fig. 2F)**. Interestingly, when hISCs were exposed to a longer 48-hour hypoxia duration, the OFE was indistinguishable from normoxia controls suggesting hypoxia-induced survival mechanisms were invoked in hISCs after 24 hours of hypoxia **(Fig. 2G)**.

To evaluate the time-dependent gene expression changes occurring in response to hypoxia and HIF1α accumulation, RNA-sequencing was performed on proliferative hISCs cultured under normoxia and severe hypoxia for 6, 24, and 48 hours **(Fig. 2H)**. Principal Component Analysis (PCA) demonstrated high agreement between technical replicates (n = 3) for each timepoint and revealed global transcriptomic changes in hISCs between normoxic and hypoxic samples. The 6-hour and 48-hour normoxic samples clustered closely on the first Principal Component (PC), whereas the 24-hour normoxic samples deviated from the other normoxic samples on the first PC. Interestingly, the 48-hour normoxic samples deviated from the 6-hour normoxic samples on the second axis, which is consistent with our previous data that hISCs mount a transient HIF-response at 24 hours following media change that resolves after 48 hours in culture (**Fig 2A,B**). These results suggest that normoxic hISCs experience a minor hypoxic event following media change that results in a subtle, but meaningful, shift in gene expression profile in line with a hypoxic response and that following this shift, the hISCs return to a slightly altered gene expression profile.

For hypoxic samples, the 6-hour samples clustered closest to the 24-hour normoxic samples, which is consistent with the previous observation that the 24-hour normoxic hISCs were undergoing a minor HIF-response. By contrast, the 24-hour and 48-hour hypoxic samples clustered together but deviated significantly from all other samples on the first PC, indicating the onset of a strong HIF-response in hISCs at 24 hours of severe hypoxia that persists through 48 hours of hypoxia. The transcriptomic shift at 24 hours of hypoxia is aligned with the robust HIF1α accumulation and significant decrease in OFE after 24 hours of hypoxia, reinforcing the concept that hISCs tolerate shorter periods of hypoxia through at least 6 hours but require transcriptional changes at 24 hours to regain organoid forming activity at 48 hours **(Fig. 2F,G)**.

To determine if durations of severe hypoxia promoted differentiation, transcriptomic analysis was performed to evaluate the expression of hallmark lineage markers for hISCs and main classes of differentiated cells **(Fig 2I)**.^47,48^ Consistent with decreased OFE at 24 hours of hypoxia, hISCs show decreased expression of stem and progenitor marker genes. Interestingly, this decreased expression continues through 48 hours of hypoxia, even though OFE is restored to normoxic levels, which is consistent with the changes seen in our PCA (**Fig 2H**). The data show very low expression and no appreciable trends in hypoxia-associated changes of genetic markers for Absorptive Enterocytes (ALPI, SI) and secretory lineages, Paneth-(DEFA5), Enteroendocrine-(CHGA, CHGB), Tuft-(PTGS1), Goblet-(MUC2), and BEST4 cells (BEST4) compared to markers of proliferative hISCs and progenitor cells. These data indicate that hypoxia alone does not promote differentiation of hISCs.

### Hypoxia primes an hISC transcriptional state to respond to extrinsic interleukin signals and intrinsic downstream mediators of inflammation

To investigate the potential molecular and physiological context associated with these time-dependent transcriptomic changes, Gene Set Enrichment Analysis (GSEA) **(Fig. 3A)** was used to evaluate the comprehensive gene expression changes observed in our RNA sequencing data. GSEA demonstrates that the hypoxia-related gene set is highly enriched at all timepoints compared to controls, providing confidence in the pathway-based analysis tools^49,50^ GSEA findings indicate hISC monolayers regulate a subset of pathways consistently across all hypoxia durations while other pathways are regulated in a dynamic manner. Specifically, transcriptomic analyses on hISC monolayers revealed enriched gene sets for inflammatory signaling through specific interleukin-mediated pathways, (e.g., Inflammatory Response, IL6 JAK STAT3 Signaling, IL2 STAT5 Signaling, and INFα/γ Responses) (**Fig. 3A**).

**Figure 3.**
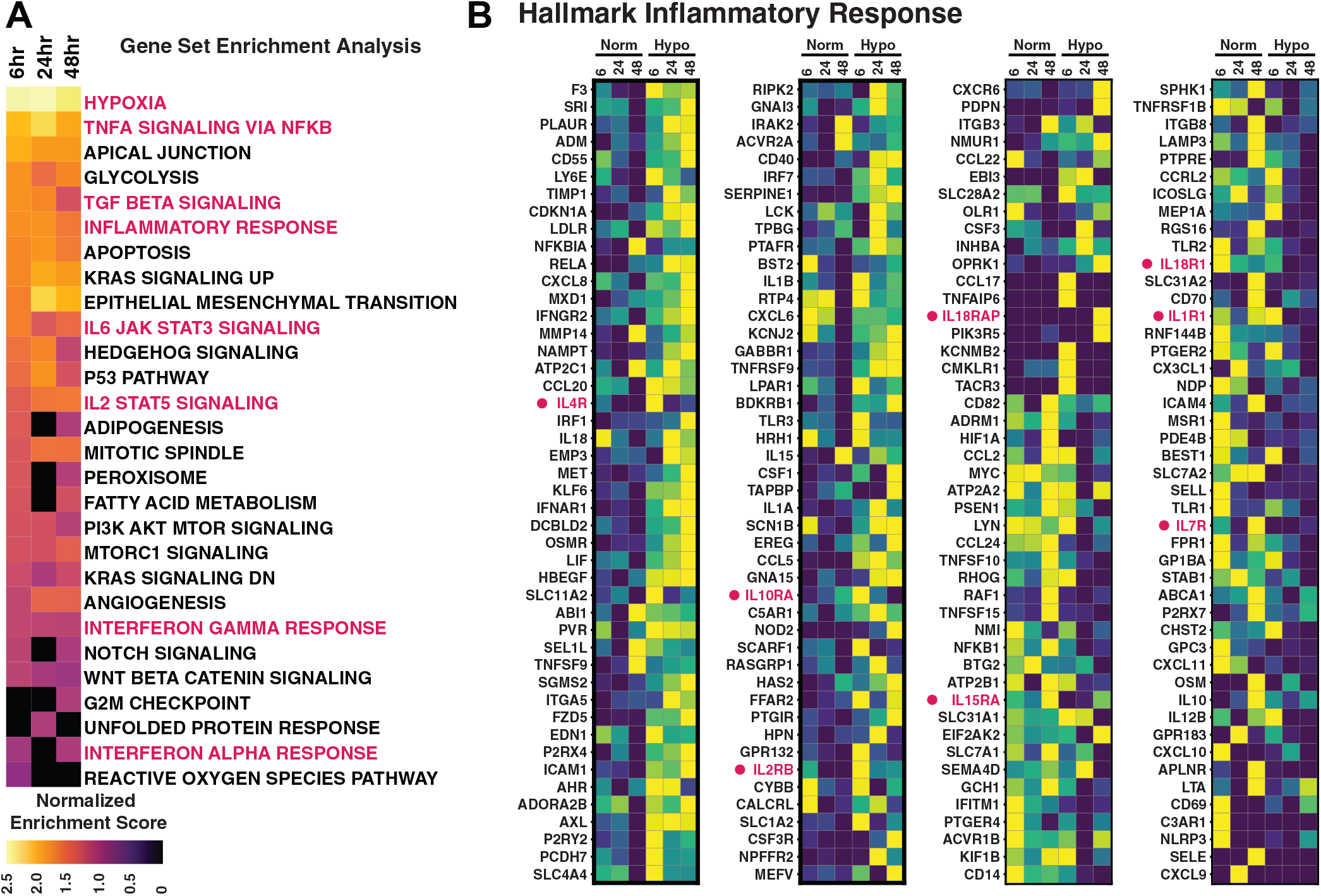
Characterizing the transcriptomic response to different durations of hypoxia by primary human intestinal epithelium. **A)** Hallmark GSEA results for differentially expressed genes at each time point of hypoxia shown compared to normoxic controls. DEGs from RNAseq from each timepoint from **2H** were compared using GSEA and ranked by Normalized Enrichment Score. Gene sets related to hypoxia or inflammatory signaling are highlighted in pink. **B**) Heatmaps of gene expression from bulk RNAseq data from **2H** for all genes from the Hallmark Inflammatory Response GSEA gene expressed in the bulk data set. Interleukin receptors from the Hallmark Inflammatory Response gene set that are expressed in our dataset are highlighted in pink.

Within the gene set for ‘Hallmark Inflammatory Response’ ∼4.5% of the genes (8/184) were interleukin (IL) receptors and all demonstrated some form of hypoxia-dependent regulation, thus we focused our analysis on all IL receptors^4,10,14^ with the goal of revealing how hypoxia-related inflammatory conditions might extrinsically signal to hISCs **(Fig. 3B)**. Interleukins and their cognate receptors have well established roles in regulating stem cell proliferation in diverse tissue types^51-60^. Of 45 IL receptors, 91% had detectable expression across all durations of hypoxia. Differential gene expression (DGE) between normoxic and hypoxic environments revealed a subset of 13 IL receptors (IL1R1, IL1R2, IL1RAP, IL1RAPL1, IL2RG, IL4R, IL11RA, IL13RA1, IL17RA, IL17RC, IL17RE, IL18R1, IL20RA) that were regulated by hypoxia in a time-dependent fashion, potentially indicating specific roles during early (6 hours), mid (24 hours), and late (48 hours) durations of severe hypoxia.

To compare hypoxia-dependent changes in IL receptor expression, we looked at fold change differences in gene expression for the 45 IL receptor genes observed in our dataset. IL1RAPL1, IL1R2, IL10RB, IL22RA1, IL6R, IL11RA, IL17RE, IL1RN, IL2RG, IL4R, IL17RA, and IL17RB were considered early responders, as all showed hypoxia-dependent up-or down-regulated expression at 6 hours (**Fig 4A**). Of the early responders IL1RN, IL6R, IL10RB, IL17RB, and IL22RA1 were considered early persistent responders as their hypoxia-dependent expression changes were consistent through 24 and 48 hours of hypoxia **(Fig 4A-C)**. IL1RAP, IL1R1, IL18R1, and IL20RA were considered mid-responders as their regulation patterns changed beginning at 24 hours of hypoxia. Of these, only IL1RAP had significant change in regulation through 48 hours. IL13RA1 and IL17RC were considered late responders as their hypoxia-dependent gene expression patterns were only observed following 48 hours of hypoxia. IL1R2 and IL11RA were considered dynamic responders as their expression patterns changed at 6 and 48 hours of hypoxia. The significant trends were verified by qPCR for the subset of IL receptors that were consistently upregulated (IL6R, IL10RB, IL22RA1) or downregulated (IL17RB) **(Fig. 4D)**. To confirm that hISCs expressed the IL receptors in vivo and in the MPS culture system, we compared IL receptor expression from our recent single cell transcriptomic atlas of the human intestine to single cell transcriptomic dataset for hISCs cultured as monolayers in media that supports hISC expansion **(Fig. 4E,F)**.^48,61^ The data show that all members of the subset of hypoxia-regulated IL receptors have measurable expression in hISCs both in vivo and in vitro. These results show hISC undergo time-dependent changes in IL receptor expression during acute and chronic hypoxic events that primes hISCs to respond to extrinsic inflammatory signals.

**Figure 4.**
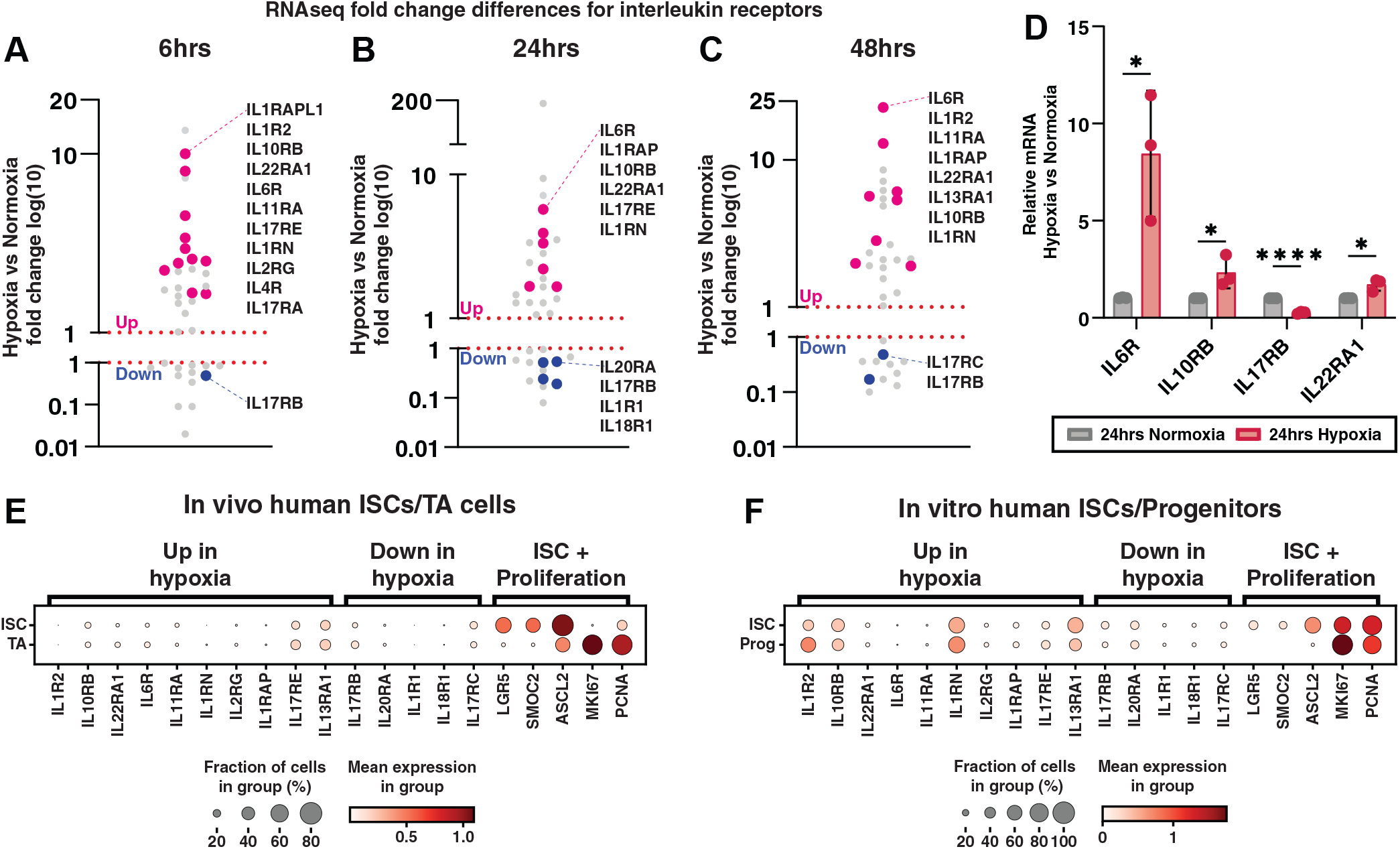
Hypoxia-induced changes in hISC gene expression for interleukin receptors. **A-C**). Fold change comparison of expressed IL receptor genes from each duration of hypoxia in **2H** is shown. Differentially expressed genes that are upregulated in hypoxia compared to their normoxic counter parts are shown above 1 and genes downregulated in hypoxia are shown below 1. Pink dots signify genes that are significantly upregulated in hypoxia at a given timepoint and blue dots represent genes that are significantly downregulated in hypoxia at a given time point. Grey dots represent IL receptor genes that are not significantly differentially expressed at a given time point. Text lists genes represented by the colored dots sorted by descending fold change. **D**) qRT-PCR measured after 24 hours of normoxia or hypoxia for IL receptor genes that are consistently up- or down-regulated at either 6, 24, or 48 hours of hypoxia. **E**) Dot plot showing DEG interleukin receptors from (**A-C**) and hISC and proliferation marker genes from hISC and TA cells from scRNAseq data of primary human intestinal epithelium. **F**) Dot plot showing DEG interleukin receptors from (**A-C**) and hISC and proliferation marker genes from hISC and TA cells from scRNAseq data of in vitro primary hISC and progenitors.

### Hypoxia increases proportion of hISCs residing in G1 phase

As a baseline, in the absence of interleukins, we determined the impact of hypoxia on regulating proliferation and cell cycle phases **(Fig 5A)**. hISC monolayers were pulsed with EdU for the last 2 hours of culture to quantify cells in S-phase during normoxia or hypoxia, then co-stained for general cell-cycle marker KI67 (**Fig 5B**). Individual nuclei were segmented using Cellpose and the percent of cells in S-phase (EdU^+^) or in any cell cycle phase (KI67^+^) were quantified.^62,63^ Confocal images were used to train the algorithm for precise nuclear segmentation (Bisbenzimide+). Approximately 150,000 nuclei were automatically identified (i.e., segmented) across all experimental conditions (**Fig 5B)**.^62,63^ After segmentation, KI67 and EdU channels were overlaid onto the automatically segmented nuclei to quantify the mean fluorescence intensity and percent of cells positive for each cell cycle marker on a per-nuclei and per-colony basis. The number of cells in mitosis (M-phase) were evaluated by counting condensed chromosomes that were stained by KI67 and morphologically identifiable (**Fig 5C**).

**Figure 5.**
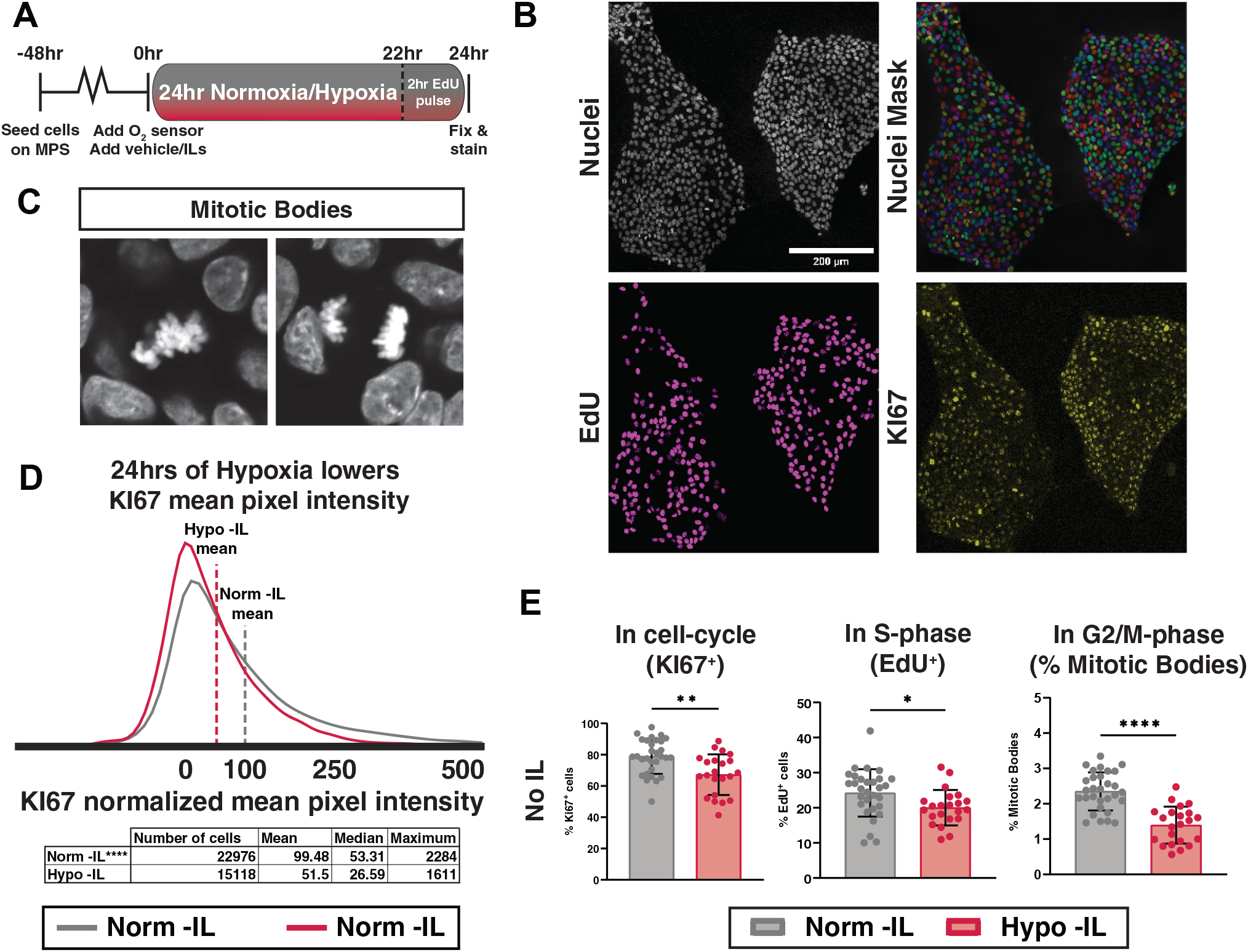
Severe hypoxia increases proportion of hISCs residing in G1 phase. **A**) Experimental schematic to assess impact of hypoxia and interleukins on cell cycle progression in human intestinal epithelial cells. **B**) Example of raw confocal imaging data showing Bisbenzimide+ nuclei (top), Cellpose segmented nuclei mask, (second from top), EdU (second from bottom), and KI67 (bottom). **C**) Example of mitotic bodies found in KI67 channel. **D**) Histogram showing distribution of normalized KI67 mean pixel intensities across Normoxic and Hypoxic conditions without interleukin treatments. Table shows key descriptive statistics that highlight statistically significant differences in distributions. **** p<0.0001 by Kolgorov-Smirnov test of cumulative distributions. **E**) Percent of KI67^+^, EdU^+^, and Mitotic Bodies counted per colony in normoxia and hypoxia conditions without treatment from interleukins. Dots represent per-colony averages of single-cell mean-pixel intensity values from separate normoxic or hypoxic colonies.

To determine whether hypoxia caused a specific arrest or delay in any particular cell cycle phase, we leveraged a recent study which shows that KI67 expression levels can reliably predict cell cycle phases.^64^ KI67 levels can serve as a marker for cell cycle progression and exit, with the lowest levels of KI67 in G1/G0 phase, intermediate levels during S-phase, and highest levels in G2/M phase^64^. We evaluated the levels of KI67 in hISCs for segmented nuclei from ∼38,000 cells and found that severe hypoxia significantly decreased KI67 under severe hypoxia (**Fig. 5D)**. Hypoxic hISC cultures showed approximately 67.2% of cells undergoing cell cycling (KI67^+^), a significant decrease compared normoxic conditions where 78.9% of cells reside in the cell cycle (**Fig 5E**).^64^ More granular analysis revealed reductions in the number of cells in S-phase (EdU^+^) and M-phase (mitotic bodies) (**Fig 5E**). The lower levels of KI67, combined with the significant decrease in S- and M-phase cells, suggests that 24 hours of severe hypoxia likely causes either G1 arrest or lengthening of G1. Transcriptomic data does not show increases in differentiated lineage markers (**Fig. 2I)** indicating that hypoxia is not pushing hISCs to differentiate as they exit the cell cycle. Together, these findings demonstrate that over 24 hours of hypoxia, hISCs negatively regulate progression through the cell cycle and that hypoxia likely promotes G1 dormancy in hISCs.

### Hypoxia alters hISC cell cycle regulation by specific interleukins

We next explored whether the hypoxia-regulated IL receptors could impact hISC cell cycle when in the presence of their cognate interleukins: IL1α, IL1β, IL2, IL4, IL6, IL10, IL13, IL17^A/F^ (heterodimer), IL22, and IL25. hISC monolayers were treated with ILs under normoxia or hypoxia for 24 hours, to simulate ILs that might be present during an chronic inflammatory response, before fixing, staining, and imaging^8,12,65-68^. No ILs caused significant impacts on the total proportion of proliferating cells (KI67^+^) after 24 hours of hypoxia; however, at 24 hours of normoxia or hypoxia, hISCs shift cell cycle phase dynamics in response to interleukin signals (**Fig 6A-F**). IL1α increases the proportion of hISCs in S-phase in normoxia, but this effect is largely abrogated in hypoxic hISCs (**Fig 6A**). IL2, IL4, and IL25 significantly increase the percentage of hISCs in M-phase only in hypoxia, with no change noted in normoxia (**Fig 6B-E**); whereas IL22 affects normoxic hISCs, but not hypoxic (**Fig 6F**). IL6, IL10, IL13, and IL17^A/F^ had no effects on cell cycle phase dynamics **(Fig. 7A-D)**. The IL1α -dependent decrease in S-phase cells with no increase in M-phase hISCs suggests that IL1α further promotes G1 arrest or elongation under hypoxia, thereby reducing observable S-phase cells. IL2, IL4, and IL25-dependent increases in M-phase hISCs with no observable decrease in KI67^+^ cells suggest slow progression of hISCs through M-phase under hypoxic conditions. These indicate that hypoxia primes hISCs to respond to ILs differently then hISCs in normoxia

**Figure 6.**
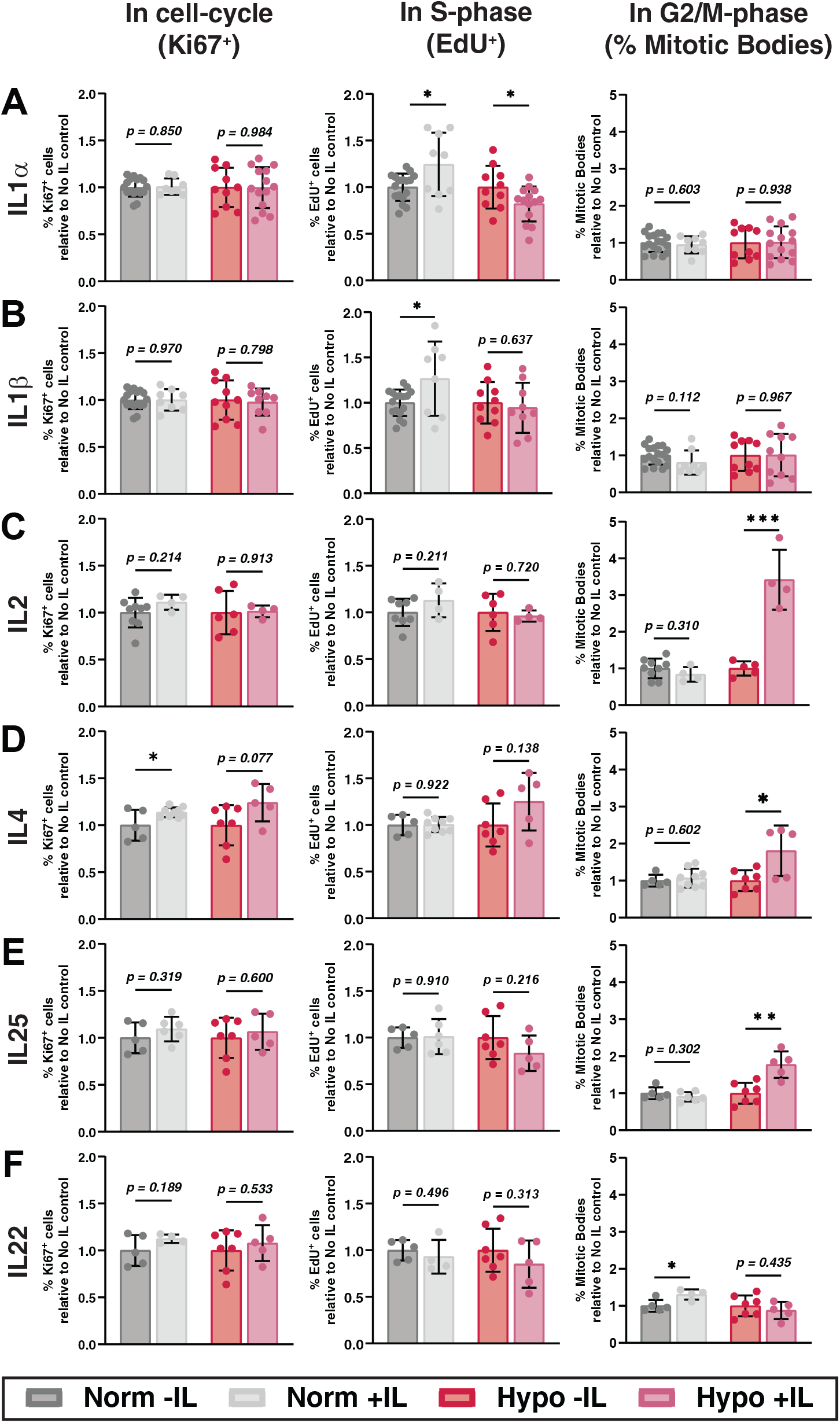
Severe hypoxia primes hISCs to respond differently to regulation of cell cycle by interleukins. **A**) Quantification of cell cycle progression metrics from **5E** for hISCs treated with 24 hours of normoxia or hypoxia with or without IL1α. **B**) Quantification of cell cycle progression metrics from **5E** for hISCs treated with 24 hours of normoxia or hypoxia with or without IL1β. **C**) Quantification of cell cycle progression metrics from **5E** for hISCs treated with 24 hours of normoxia or hypoxia with or without IL2. **D**) Quantification of cell cycle progression metrics from **5E** for hISCs treated with 24 hours of normoxia or hypoxia with or without IL4. **E**) Quantification of cell cycle progression metrics from **5E** for hISCs treated with 24 hours of normoxia or hypoxia with or without IL25. **F**) Quantification of cell cycle progression metrics from5**4E** for hISCs treated with 24 hours of normoxia or hypoxia with or without IL22.

**Figure 7.**
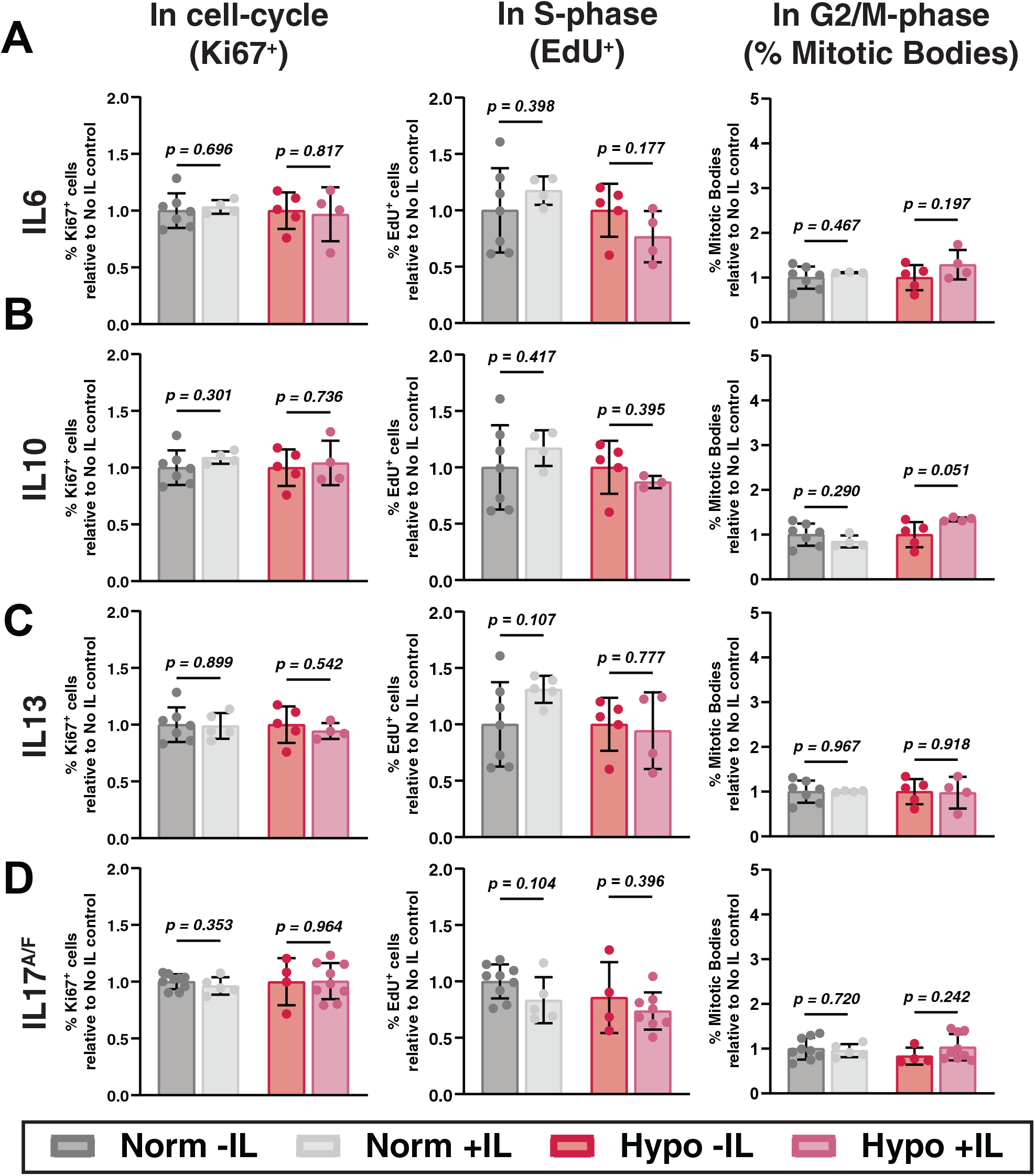
Interleukins with no significant effect on hISCs regardless of oxygen condition. **A**) Quantification of cell cycle progression metrics from **5E** for hISCs treated with 24 hours of normoxia or hypoxia with or without IL6. **B**) Quantification of cell cycle progression metrics from **5E** for hISCs treated with 24 hours of normoxia or hypoxia with or without IL10. **C**) Quantification of cell cycle progression metrics from **5E** for hISCs treated with 24 hours of normoxia or hypoxia with or without IL13. **D**) Quantification of cell cycle progression metrics from **5E** for hISCs treated with 24 hours of normoxia or hypoxia with or without IL17^A/F^.

### Hypoxia affects hISC activity in response to interleukin treatment

While the previous experiments were designed to focus on the impact of hypoxia-dependent IL receptor priming on proliferation dynamics, we next sought to probe whether this IL receptor priming would regulate stem cell activity in the presence of cognate interleukins. hISC monolayers were exposed to 24 hours of normoxia or severe hypoxia, then dissociated to single hISCs and plated in conventional 3D organoid assays to evaluate OFE of clonal hISCs in the absence or presences of ILs (**Fig 8A**). Similar to OFE assays in the CellRaft arrays (**Fig 2F**), we found that hypoxia induced a decrease in OFE at 24 hours (**Fig. 8B,C**). IL6, IL10, and IL13 (**Fig 8D-F**) increased OFE in a hypoxia-independent manner, while IL1β, IL2, IL4, and IL25 demonstrated hypoxia-dependent increases in OFE (**Fig 8G-J**), thereby rescuing the negative impacts of hypoxia on hISC. No effects on OFE were observed for IL1α, IL17^A/F^, and IL22 (**Fig 8K-M**).Together, these findings show 24 hours of hypoxia primes hISCs to respond to cognate ILs differently in hypoxia than in normoxia by altering cell cycle phase dynamics and in some cases enhancing hISC activity.

**Figure 8.**
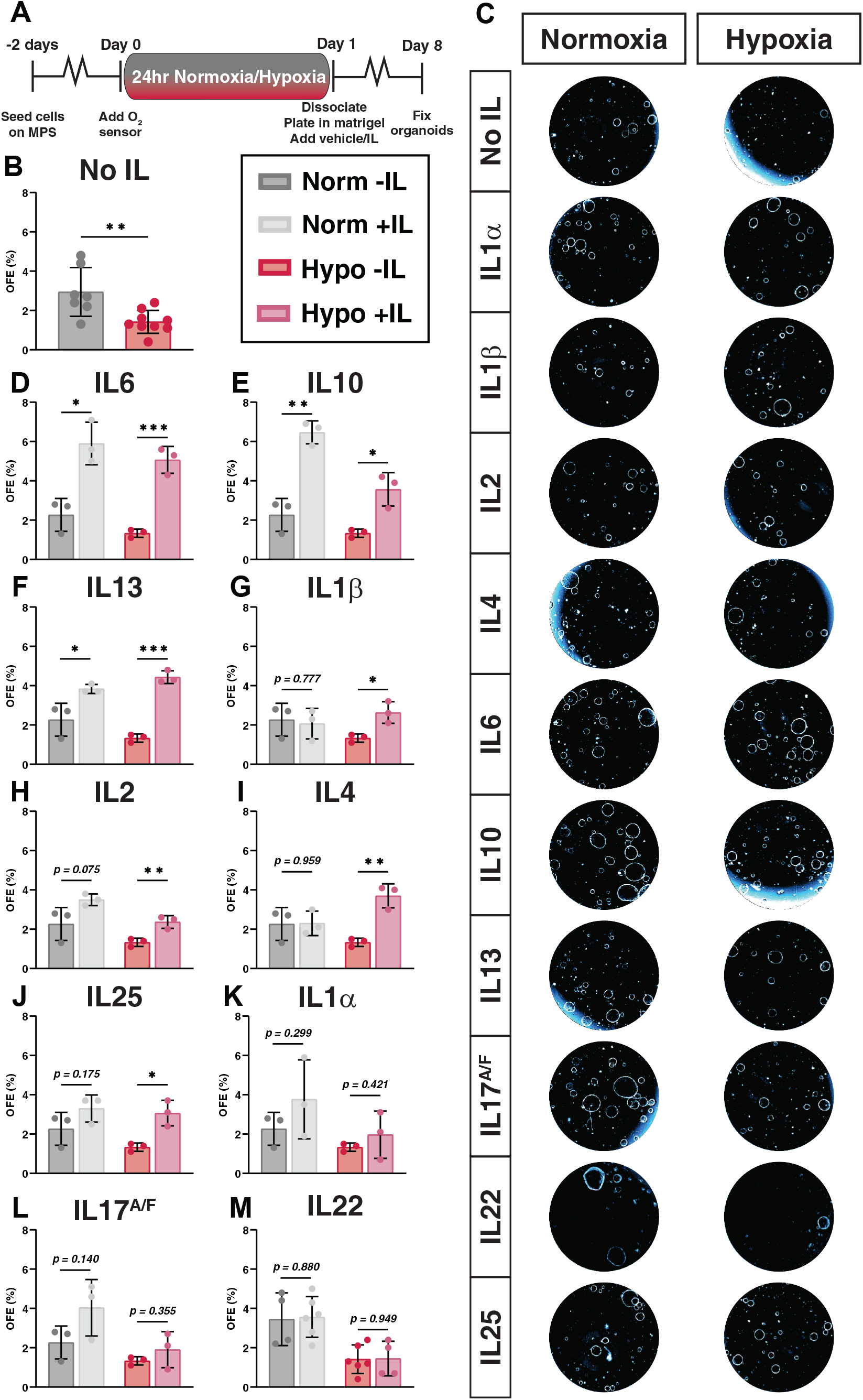
hISCs demonstrate hypoxia-dependent rescue of organoid forming efficiency when treated with specific interleukins. **A**) Experimental schematic to assess impact of hypoxia, then reoxygenation and interleukins on functional stem cell activity. **B**) Impact of hypoxia and reoxygenation on functional stem cell activity without treatment of interleukins. **C**) Representative images of organoids from conditions in **B,D-M**. Images have been pseudocolored to highlight outer membranes of organoids. **D**) Impact of hypoxia and reoxygenation plus IL6 from on functional stem cell activity. **E**) Impact of hypoxia and reoxygenation plus IL10 from on functional stem cell activity. **F**) Impact of hypoxia and reoxygenation plus IL13 from on functional stem cell activity. **G**) Impact of hypoxia and reoxygenation plus IL1β from on functional stem cell activity. **H**) Impact of hypoxia and reoxygenation plus IL2 from on functional stem cell activity. **I**) Impact of hypoxia and reoxygenation plus IL4 from on functional stem cell activity. **J**) Impact of hypoxia and reoxygenation plus IL25 from on functional stem cell activity. **K**) Impact of hypoxia and reoxygenation plus IL1α from on functional stem cell activity. **L**) Impact of hypoxia and reoxygenation plus IL17^A/F^ from on functional stem cell activity. **M**) Impact of hypoxia and reoxygenation plus IL22 from on functional stem cell activity.

## DISCUSSION

Human health conditions that cause loss of oxygen to the intestinal epithelium result in rapid loss of the differentiated cells, requiring quick regeneration from the stem cell pools located at the base of the crypts^6,69,70^. Evaluating the impact of hypoxia on hISCs in vivo is challenging due to ethical considerations and the inability to precisely control many experimental variables. While in vitro systems are available to regulate ambient oxygen levels, these platforms are limited in that oxygen levels are monitored in large volume incubators and not localized to the cell-medium interface. Here we developed a flexible and oxygen-tunable MPS that addresses many limitations of conventional systems and represents a next-generation platform to study hypoxic injury across a broad range of cell and tissue types. While not explored in this study, the flexible design and small scale of the MPS will enable co-culture of hISC with other cell types, such as immune cells, which are key players in hypoxia-related intestinal conditions.

Murine intestinal epithelial monolayers were shown to experience repeated bouts of hypoxia during conventional cell culture methods.^39^ Upon media changes, oxygen levels were at saturation but the levels substantially reduced in media over time, ostensibly by cellular metabolism, thereby creating a hypoxic environment and inducing a HIF1α response.^39^ These observations indicate that oxygen diffusion from ambient incubator gas mixes was insufficient to compensate for the oxygen metabolism of mouse intestinal epithelial cells submerged in culture media.^39^ Using our newly developed MPS that supports 2D cultures of hISCs, we show similar findings in that fresh media at the cell surface interface is saturated at about ∼12% O_2_ from atmospheric gasses and that hISCs metabolize O_2_ levels in the media from ∼12% to ∼3% in approximately 3 hours. At that point, O_2_ levels stabilize but are sufficiently low to induce a small but appreciable hypoxia-induced HIF1α response after 24 hours of submersion culture. Our findings challenge general assumptions regarding oxygen availability following regular medium changes in many conventional cell culture systems. More specifically, our data suggest that any cells cultured in submersion culture can experience unappreciated HIF1α-dependent transcriptional changes following media changes, confounding interpretations.

Our findings show that hISCs are highly resilient to severe hypoxia for up to 48 hours. There was a decrease in functional hISC activity at 24 hours of hypoxia, but this resolved at 48 hours with no significant difference in OFE compared to normoxic controls. Several studies in rodent and human intestines highlight the role of endoplasmic reticulum (ER) stress and the unfolded-protein response (UPR) during acute hypoxic episodes.^39,71,72^ The UPR is considered to mediate cell death by activating the intrinsic apoptotic cascade; however, several more contemporary reports in the context of cancer cell lines indicate that unresolved ER-stress, which can occur during long durations of hypoxia^73^, can induce necroptosis.^73-75^ We found no clear evidence of increased apoptosis during 48 hours of severe hypoxia, and while necroptosis pathways were not evaluated, there was no decrease in hISC activity at 48 hours of hypoxia, strongly supporting that other cell death pathways are being repressed in hISCs through 48 hours of hypoxia.

Transcriptomic profiling of hISCs during chronic and severe hypoxia revealed enrichment of regulated genes related to inflammation and inflammation responses. Among these genes we noticed many cytokine/interleukin receptors were significantly regulated in a hypoxia-dependent manner. While the interleukin responses mediated through these receptors impact many different aspects of cell biology, we focused our analysis on how hypoxia-induced changes in hISC expression of IL receptors, could impact cell cycle dynamics and stem cell activity through stimulation using their cognate ILs. In the absence of ILs, the baseline hISC response to 24 hours of hypoxia was a significant decrease in the overall number of cycling cells (KI67^+^) with notable reductions in numbers of cells in each phase of the cell cycle. While the number of cycling cells was reduced at 24 hours of hypoxia based on KI67, the lack of differentiated cell markers and absence of reduced OFE of hISCs at 48 hours suggests hISCs do not exit cell cycle or differentiate as the result of severe hypoxia but rather enter a ‘safe mode’ characterized by a reversable dormant state, reminiscent of a reserve ISC state^76^. Further genetic and molecular analysis will be necessary to reveal these mechanisms.

During inflammatory hypoxia, local immune cells express a complex mixture of ILs near hISCs.^8^ In our model, hypoxia primed hISCs to respond to subsets of ILs by regulating cognate IL receptors. Stimulation of hypoxia-regulated IL receptors by their cognate ILs produced notable hypoxia-dependent increases in G2/M-phase cells for IL2, IL4, and IL25 that generally rescued the baseline hypoxia-dependent loss of G2/M-phase cells. Less notable, IL1α caused a modest but significant reduction in S-phase cells. There is a collection of studies describing how IL2 and IL4 promote proliferation in immune cells^53,56,77,78^, and studies in colon and pancreatic cancer cells demonstrate that IL4 enhances proliferation^79,80^, but we found no studies describing the role of these ILs on mouse or human ISCs. IL25 is expressed by tuft cells during parasitic worm infections.^81-84^ Secreted IL25 interacts with the cognate heterodimer receptors, IL17RA and IL17RB^85^, on Innate Lymphoid Cells Type 2 (ILC2s), which signal back through IL-4 and IL-13 to the epithelium to aid worm clearance.^81,83,84,86^ IL17RA/B expression has not previously been reported in hISCs, suggesting a novel role for IL25 signaling in hISCs.

Under 24 hours of hypoxia, hISCs demonstrate less active progression through the cell cycle and lower organoid forming efficiency. hISCs show increased stem cell activity in response to IL6, IL10, and IL13, regardless of hypoxic conditioning. On the other hand, hypoxia induces hISCs to respond to IL1β, IL2, IL4, and IL25 by increasing stem cell activity in a hypoxia-dependent fashion. IL2 and IL4 promote cell survival effects by stimulating the Akt-pathway^87^, and although there are fewer clear links between IL25 (IL17RA/B) and cell survival, the IL17RA/B receptors can stimulate NF-kB activation^88,89^, which has cell survival effects under some stress stimuli.^90^ While hypoxia alone can stimulate NF-kB activation through HIF1α^8,91^, IL25 could amplify NF-kB activation to increase hISC survival during severe and prolonged hypoxic episodes. Interestingly, IL2, IL4, and IL25 stimulated increases in G2/M in hISCs under hypoxia, with no change seen in hISCs under normoxia. G2/M arrest or delay by these three cytokines might protect cells from undergoing checkpoint-induced cell death as the completion of cytokinesis could be compromised by the lack of energetic and metabolic resources under severe hypoxia.^92^ While many ILs have cell cycle effects on hISCs under normoxia, we see that many of these effects change and produce novel responses that are only seen in the context of hypoxia. As such, it appears hypoxia-primed hISCs likely respond differently to immune stimuli than normoxic hISCs, laying the foundation for studies into unique, hypoxia-dependent genetic and biochemical mechanisms, which preserve and restore hISC activity during and after hypoxic injury.

We acknowledge that deconstruction of the complex microenvironment of the in vivo human stem cell niche limits some physiological interpretations. However, second generation studies using our MPS can build on the findings of this study to test complex mixtures of hypoxia-related cytokines, interactions of multiple cell types, and tunable O_2_ concentrations to more accurately model and study hISC behaviors under inflammatory hypoxia, more acute forms of oxygen deprivation, and clinically relevant reperfusion. Previous iterations of the MPS have shown the ability of the MPS to dynamically ‘reperfuse’ hypoxic cultures with O_2_, enabling studies of other modes cell death, such as ferroptosis, which likely have a role in reperfusion injury due to reactive oxygen species production^26,93,94^, or pyroptosis, which are relevant to inflammasome-mediated cell death and are complex to model as they involve microbes and immune cell components.^95,96^ As such the MPS is currently being optimized for co-culture of hISCs, immune cells, and anaerobic commensal or pathological microbiota with the goal of creating a more physiologically relevant environment to explore many facets that impact hISC function and behavior.

## MATERIALS AND METHODS

### Primary human crypt isolation and intestinal epithelial stem cell culture

A surgical specimen of human small intestine (jejunum) was obtained from a donor at UNC Hospitals with consent of the patient (under the approved protocol UNC IRB #14-2013). Villi and crypts were detached from the specimen by incubation in a chelating buffer for 75 min at 20°C followed by vigorous shaking in a 50 mL conical tube. The chelating buffer was composed of EDTA (2 mM), dithiothreitol (DTT, 0.5 mM, freshly added), Na_2_HPO_4_ (5.6 mM), KH_2_PO_4_ (8.0 mM), NaCl (96.2 mM), KCl (1.6 mM), sucrose (43.4 mM), D-sorbitol (54.9 mM), pH 1/4 7.4.^97^ Released crypts were expanded as a monolayer on a neutralized collagen hydrogel as described previously.^25^ Briefly, crypts were placed on the top of 1.0 mg/mL collagen hydrogels (1 ml into each well of 6-well plate (T1006; Denville, Holliston, MA)) at a density of 5,000 crypts/ well and overlaid with 4 ml of Expansion Media (EM) containing 10 mmol/L Y-27632 (S1049; SelleckChem). EM contains a mixture of advanced Dulbecco’s modified Eagle medium/F12 medium (12634010; ThermoFisher) and Wnt-3A, R-spondin 3, noggin (WRN) conditioned medium (WRN medium prepared in lab from L-WRN cells (CRL-3276; ATCC) following a published protocol^98^ at a volumetric ratio of 1:1, and supplemented with GlutaMAX (35050061; ThermoFisher), B27 supplement without vitamin A (12587010; ThermoFisher), 10 mM HEPES (15630-080; ThermoFisher), 1.25 mM N-acetyl cysteine (194603; MP Bio, Santa Ana, CA), 10 mM nicotinamide (N0636; Sigma-Aldrich), 50 ng/mL epidermal growth factor (315-09; Peprotech), 2.0 nM gastrin (AS-64149; Anaspec), 10 nM prostaglandin E2 (14010; Cayman Chemicals), 3.0 μM SB202190 (S1077; Selleckchem), 100 U/mL penicillin-streptomycin (15140122; ThermoFisher), and 50 mg/mL primocin (ant-pm-1; InvivoGen, San Diego, CA). EM was used to expand the epithelial cell numbers as monolayers or organoids. Y-27632 was present only in the first 48 hours of cell culture and was not added to subsequent media changes. The medium was changed every 48 hours. When the cell coverage was greater than 80% (typically 5 to 7 days), the epithelium was dissociated to fragments to seed onto the intestinal MPS.

### Fabrication of primary human intestinal microphysiological system (MPS)

The human intestinal MPS was fabricated from polymethylmethacrylate (PMMA) and photocured resin (Formlabs, Inc.). PMMA provided an optically transparent material with a low oxygen diffusion coefficient that could be sterilized and reused.^99^ The photocured resin provided a 3D printable bottom frame to support the PMMA device and simultaneously house the optical reader for oxygen measurements. The cell culture chamber and gas exchange channels were fabricated from 5.8 mm and 1.5 mm-thick PMMA sheets (44352, 44292; US Plastics). The microfluidic culture region was composed of 5 rectangular wells, with dimensions of 11.6 mm by 7.0 mm each. The rectangular gas channel on top of the wells was 70 mm by 45 mm by 1.5 mm. The bottom and top pieces of PMMA were laser cut from a 1.5 mm-thick PMMA sheet, while the middle piece, for the cell culture wells, was laser cut from a 5.8 mm-thick PMMA sheet. Briefly, to remove dust and burr material with minimal cracking, each PMMA surface was quickly wiped with 100% IPA solution and air-dried. All 3 pieces were pressed together between two sheets of brass using a pneumatic heat press (SwingPress 10-0403; Across International). Annealing was performed with the heat press set at 100°C and 300 psi for 2 hours, then the bonded device was cooled for 3 hours at room temperature. A rubber gasket was laser cut and placed on top of the PMMA layer to seal the device, prior to bolting together. The device was tested for leaks using red and blue dyed water and cracks were sealed by application of dichloromethane (DCM) to the seams.

In the completed device, barbed connectors screwed into the photopolymer resin top lid provide gas flow into and out of the device. A rubber gasket seals gas flow into and out of the PMMA gas exchange channel. The gas exchange channel frame is bonded to the PMMA culture chamber and PMMA base support. A photopolymer resin bottom frame supports the entire PMMA device, and screws attach the top lid to the bottom frame to seal the MPS closed. The optics front-end connects via a μUSB-to-HDMI cord to a phosphorescence-lifetime fluorimeter detector and interrogates the middle culture well, where the iPOB is located.

A neutralized collagen hydrogel (2.0 mg/ml) was cast into each well of a prefabricated PMMA device and allowed to polymerize at 37°C for 1 hour. 250 μL of 1X DPBS was overlaid on the polymerized hydrogel and allowed to pre-swell for at least 5 hours at room temperature. DPBS was removed and the hydrogel was rinsed three times before overlaying with 250 μL of EM containing hISCs that had been mechanically dissociated by pipetting 10 times until cells were ∼1-10 cell-clumps. Once the hISCs formed a confluent monolayer, an iPOB was added to the epithelium to measure oxygen.

### Real-time monitoring and control of oxygen concentration in 3D culture

Oxygen concentration in the system was continuously measured with an integrated phosphorescent oxygen sensor (iPOB) using near infrared (NIR) phosphorescence lifetime fluorimetry. The iPOB is composed of porous poly(2-hydroxyethyl methacrylate) (pHEMA) gel functionalized with palladium-benzoporphyrin derivatives (Pd-BPD) that respond to local oxygen concentrations via phosphorescence quenching.^27^ The photoluminescence excitation and detection wavelengths are 630 nm and 800 nm, respectively. The iPOB (Profusa, Inc., San Francisco, CA) has been manufactured in a variety of sizes, but for all experiments a disk-shaped, 5 mm diameter and 0.5 mm thickness, iPOB was used. The optics front-end of the NIR phosphorescence lifetime fluoroscope (Profusa, Inc., San Francisco, CA) was inserted into the support frame below the MPS. The oxygen concentration was controlled via a PMMA gas-mixing microfluidic chip with tubing connecting outlets from the gas-mixing chip to the inlets of each MPS (**Fig 9A,B**). The gas-mixing chip was previously used to generate 8 concentrations of mixed gas, ranging from less than 3 μM of oxygen to 180 μM. **(Fig 9C,D**)^28^ Gas flow to the gas-mixing chip was regulated using an air flow control valve (62005K313; McMaster-Carr) and monitored with a mass flow meter with digital output (GFMS-010061; Aalborg GFM) (**Fig 9B, Left**). Hydrated mixed gas exiting the gas-mixing chip was introduced into individual MPS at a rate of 5 mL/min to prevent media evaporation. Within 30 minutes, the gas mixture equilibrated with the local MPS environment to generate the desired oxygen concentration at the hydrogel surface-media interface where oxygen was measured using the iPOB (**Fig. 1D**). The intestinal epithelium inside MPS was cultured in the mixed gas environment and compared to intestinal epithelium statically cultured in a normobaric incubator with an oxygen environment of 186 μM.

**Figure 9.**
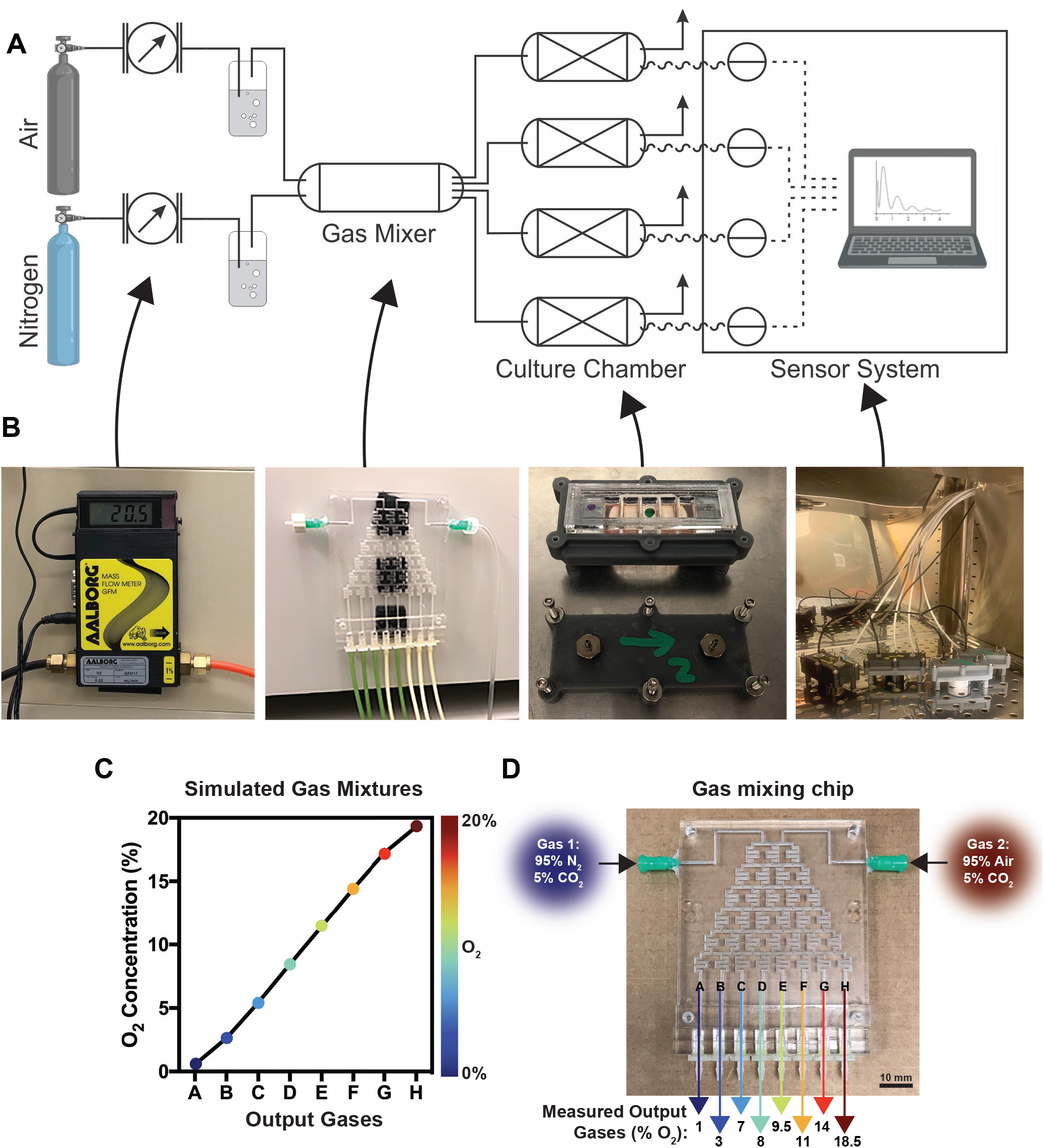
Gas mixing chip integrated into microphysiological system to generate 8 different oxygen environments. **A**) Schematic of complete microphysiological system design with mixed gases in tanks leading into flow meters. Flow meters connect into humidification 15 mL conical tube filled with sterile PBS. Humidified gases travel into each input of the gas mixer chip, and 8 concentrations of gases exit individual outlets to connect into separate microphysiological systems (devices). Device oxygen concentration is monitored using detectors that send information to a laptop located beside the incubator. **B**) Images taken of each component of the system. **C**) COMSOL Multiphysics results for the simulated gas mixtures from each outlet of the gas mixing chip. **D**) Image of the gas mixing chip and measured output oxygen concentrations recorded inside individual devices.

### Immunocytochemistry

To assess the impact of various durations and magnitudes of hypoxia on hISC proliferation, primary human intestinal epithelium and single cells isolated from each MPS chamber were stained for Ki-67. For Ki-67 staining of intestinal epithelium following hypoxia, the intestinal MPS was opened, and media was removed from each chamber. Intestinal epithelium on top of the collagen hydrogels were fixed with 4% PFA for 15 minutes. After fixation, samples were rinsed once with 1X PBS and overlaid with PBS. Samples were permeabilized for 15 minutes with 0.5% Triton-X 100 in 1X PBS. Samples were blocked for 30 minutes with a 3% BSA solution. After blocking, samples were stained for proliferation marker Ki-67 Alexa Fluor 647 (1:250 dilution in 3% BSA, Cat. No. 652407; BioLegend) for 1 hour and nuclei counterstain bisbenzimide (1:1000 dilution in 1X dPBS, Cat. No. 1155; Millipore Sigma) for 5 minutes at room temperature. To look at cell-cell contacts, tight junction protein Occludin (Cat. No. 13409-1-AP; Proteintech) was added for 1 hour, followed by incubator with the secondary antibody Cy3 (Cat. No. C2306; Sigma) for 2 hours. After staining, the gels were overlaid with 1X dPBS and stored at 4°C until imaging. All samples were imaged with a Olympus (Waltham, MA) IX81 microscope, using Metamorph Basic (Molecular Devices, San Jose, CA) software, and image analysis was performed with ImageJ.^100^

### Organoid forming efficiency assay and Fluorescence-activated cell sorting (FACS) of live cells

Following MPS culture, human intestinal epithelium was dissociated into single cells, sorted using a flow cytometer, and suspended in Matrigel^®^ on a quad CellRaft™ Array (CRA) platform (Cell Microsystems, Durham, NC).^34^ Briefly, each sample was retrieved and placed in a separate conical. 500 U/mL Collagenase IV (17104019; Gibco) was added to breakdown the scaffold and incubated for 10 minutes at 37ºC. After centrifugation, the cell pellet was rinsed twice in DPBS and re-suspended in 150 μL of 0.5 mmol/L EDTA in DPBS with 10 mmol/L Y-27632 and incubated for 5 minutes at 37ºC. The fragments were further dispersed by triturating 30 times using a 200 μL pipet tip. After centrifugation, the cell pellet was re-suspended in 500 μL TrypLE Express (12605-036; Gibco) with 10 mmol/L Y-27632 and incubated for 5 minutes at 37°C. The cell suspension was gently triturated 7 times using a 28.5-gauge insulin needle to further dissociate. 500 μL of EM was added to quench the reaction, and cells were pelleted and rinsed once in EM. After pelleting again, cells were re-suspended in 200 μL of EM containing 1% FBS. Immediately before FACS, APC Annexin V (1:200, 640941; BioLegend) was added to cells for live/dead discrimination. After staining, cells were rinsed in EM and filtered through 0.4 μm FACS tube top filter (352235; Corning). 5,000 Annexin V-live cells were isolated via FACS and re-suspended in 250 μL of Growth Factor Reduced Matrigel^®^ (354230; Corning). Cell-gel suspensions were plated in each chamber of the quad CRA. To cover the CRA, each array was centrifuged at 200 x g for 5 minutes. CRAs were polymerized for 20 minutes at 37°C in an incubator and then overlaid with EM containing 10 mmol/L Y-27632. Media was replaced every 2 days. To determine OFE, the CRA was scanned, and the total number of organoids was counted on day 2, day 4, and day 6. The OFE (%) was calculated as the total number of organoids created in one CRA chamber, divided by 5,000 cells and multiplied by 100. FACS and flow cytometry were performed using a SH800Z Cell Sorter (Sony Biotechnology, San Jose, CA).

### Flow cytometry

Cells collected from normoxic and hypoxic MPS were dissociated following the protocol above up to the step of addition of TrypLE Express. Following trituration with the insulin needle and quenching with the EM, cells were pelleted and re-suspended in 100 μL of 1X DPBS. Approximately 50,000 cells from each chamber were fixed by adding 400 μL of 4% PFA solution while being constantly vortexed to prevent cell aggregation. After fixation, single cells were pelleted and re-suspended in 0.3% Triton-X for 15 minutes to permeabilize the cell membrane. Single cells were pelleted and re-suspended in a staining solution of DMEM containing 1% FBS and anti-Ki-67 Alex Fluor 647 (1:250 dilution in 3% BSA, Cat. No. 652407; BioLegend) for 1 hour on ice. After staining, cells were rinsed in PBS and filtered through 0.4 μm FACS tube top filter (352235; Corning). Ki-67-positive single cells were quantified by flow cytometry.

### qRT-PCR

To assess the expression of genes that are responsive to hypoxia, human intestinal epithelial samples from each MPS were collected for qRT-PCR analysis. Briefly, cells attached to collagen hydrogels were lysed in 200 μL of RNA Lysis buffer (AM1931; ThermoFisher). Total RNA was extracted using RNAqueous-Micro Total RNA Isolation Kit (AM1931; ThermoFisher) according to manufacturer’s protocols. cDNA was generated from ∼2 ng of total RNA from each sample using iScript Reverse Transcription Supermix for qRT-PCR (170-8891; BioRad) according to manufacturer’s protocols. cDNA was diluted 1:20 and 1 μL was used for qRT-PCR using the following Taqman probes: HIF1α:Hs00153153_m1, BNIP3: Hs00969289_m1, SLC2A1 (GLUT1): Hs00892681_m1, VEGFA: Hs00900055_m1, IL6R: Hs01075664_m1, IL10RB: Hs00175123_m1, IL17RB: Hs00218889_m1, IL22RA1: Hs00222035_m1 (Applied Biosystems) and SsoAdvanced Universal Probes Supermix (1725281; BioRad) according to manufacturer’s protocols. qRT-PCR was carried out in a StepOnePlus Real Time PCR System (Applied Biosystems). For each sample and experiment, triplicates were made and normalized to 18S mRNA levels. Fold change was expressed relative to normoxic controls using ΔΔCT analysis.^101^ All statistics for gene expression were generated using a Student’s t-test. In all statistical analysis, *p* < 0.05 was considered significant.

### Bulk RNA-sequencing

To investigate the dynamic response of hISC to hypoxia at the whole transcriptomic level, we performed RNA-seq on human intestinal epithelium exposed to 6 hours, 24 hours, or 48 hours of hypoxia inside intestinal MPS, along with normoxic control intestinal MPS which were cultured for each respective time point with no hypoxia exposure. RNA samples were collected from each intestinal MPS, and Total RNA was extracted using RNAqueous-Micro Total RNA Isolation Kit (AM1931; ThermoFisher) according to manufacturer’s protocols and stored at −80°C. To assess RNA quality prior to submission for sequencing, an RNA integrity number (RIN) was measured using the Agilent 2100 Bioanalyzer. After confirmation that each sample had a RIN of at least 8, integrated fluidic circuits (IFCs) for gene expression and genotyping analysis were prepared using the Advanta™ RNA-Seq NGS Library Prep Kit for the Fluidigm Juno™.

### Informatics

Gene level expression was obtained through pseudo alignment of reads to human genome GRCh38 using Kallisto.^102^ Principal component and sample correlation analysis were done with Bioconductor packages Biobase, cluster and qvalue.^103^ Expression values for plotting were obtained by TMM normalization across all samples using EdgeR package and differential expression between normoxic and hypoxic samples at each time point was calculated from raw counts with DESeq2.^104,105^ GSEAs were performed using all hallmark gene sets on n = 3 samples from each duration of hypoxia (6-, 24-, and 48-hours) to its corresponding normoxic control separately. These comparisons were ranked by NES.^106,107^

## ABBREVIATIONS

(IBD): Inflammatory Bowel Disease
(ISC): Intestinal stem cell
(hISC): Human intestinal stem cell
(MPS): Microphysiological system
(iPOB): integrated Phosphorescent Oxygen Biosensor
(NIR): Near Infrared
(2D): 2-Dimensional
(OCLN): Occludin
(OFE): organoid forming efficiency
(CRAs): CellRaft™ Arrays
(PCA): Principal Component Analysis
(PC): Principal Component
(GSEA): Gene Set Enrichment Analysis
(IL): interleukin
(DGE): Differential gene expression
(ER): endoplasmic reticulum
(UPR): unfolded-protein response
(ILC2s): Innate Lymphoid Cells Type 2
(DTT): dithiothreitol
(EM): Expansion Media
(PMMA): polymethylmethacrylate
(DCM): dichloromethane
(pHEMA): poly(2-hydroxyethyl methacrylate)
(FACS): Fluorescence-activated cell sorting
(RIN): RNA integrity number
(IFCs): integrated fluidic circuits

## ACKNOWLEDGEMENTS

The authors would like to thank all the members of the Magness Laboratory, specifically Dr. Meryem Ok, and the Biointerface Lab, along with Drs. Michael Gamcsik and Bailey Zwarycz for helpful discussions. The authors thank the Center for Gastrointestinal Biology and Disease (CGIBD) and the CGIBD’s Advanced Analytics Core under NIH grant P30 DK034987. This work was done in collaboration with the National Science Foundation (EEC-1160483) through a NSF Nanosystems Engineering Research Center (NERC) for Advanced Self-Powered Systems of Integrated Sensors and Technologies (ASSIST), CCSS-1846911 (M.D.) and CBE/EBS-2033997 (M.D., S.T.M.).. K.R.R. was supported by the National Institutes of Health under award number F31DK118859-02, the 2019 Howard Hughes Medical Institute Gilliam Fellowship Award, F32DK124929 (J.R.B.), R01DK109559 (S.T.M.), the Katherine E. Bullard Charitable Trust for Gastrointestinal Stem Cell and Regenerative Research (S.T.M.), and CGIBD Pilot & Feasibility Award P30 DK034987 (M.D./S.T.M.). We thank Gabrielle Cannon at the Advanced Analytics core at UNC Chapel Hill.

## AUTHOR CONTRIBUTIONS

K.R.R. designed the study and conducted the experiments, analyzed data, and wrote the manuscript.

R.J.B. conducted experiments, analyzed RNA sequencing results, and wrote the manuscript

M.J.C. conducted experiments and analyzed RNA sequencing results.

J.B. conducted experiments, helped analyze data, and reviewed the manuscript

J.L. conducted experiments and analyzed data

J.M.T. conducted experiments

C.M.H. helped analyze data

K.A.B. helped analyze data

S.J. conducted experiments

V.A.P. worked with K.R.R. to build and integrate all components of the microphysiological systems.

M.Y. designed, fabricated, and validated the gas mixing chip using COMSOL Multiphysics and experimental methods.

M.D. and S.T.M. conceived the study and wrote the manuscript.

A.Z. Contributed experimental design and informatics analysis

A.T.B. Provided intellectual contributions on the topics of mechanisms of ischemia/reperfusion and hypoxic injury

